# Deep leaning auto-segmentation of tomographic datasets, repeatability and biomechanical outcomes

**DOI:** 10.1101/2025.11.06.686240

**Authors:** Felix Bowers, Richard Lane, Wahab Kawafi, Danielle Paul, Tahlia Pollock, Harry Berks, Emily J. Rayfield, Chrissy L. Hammond

**Affiliations:** Palaeobiology Research Group, School of Earth Sciences, Life Science Building, University of Bristol, BS8 1TQ; School of Translational Health Sciences, Bristol Medical School, Biomedical Sciences Building, University of Bristol, Bristol, BS8 1TD; Jean Golding Institute, Royal Fort House, University of Bristol.; School of Biological Sciences, Monash University, Monash, 3800

## Abstract

Automated segmentation of three-dimensional micro-computed tomography (CT) scan data is a critical bottleneck in computational morphometrics and biomechanical modelling across musculoskeletal biology. Although advances in imaging have generated increasingly large and complex datasets, manual segmentation remains prohibitively time-consuming, while existing deep learning solutions are often application-specific and rarely validated for their impact on downstream analyses. Here we present a generalisable computational framework for automated segmentation and biomechanical validation of skeletal structures, implemented using an attention-augmented 3D U-Net architecture. Using the adult zebrafish (Danio rerio) mandible as a representative case study, the network was trained on 47 manually segmented specimens and evaluated using a combined Dice-Hausdorff metric that integrates both volumetric and surface accuracy to capture biologically relevant morphology better than Dice score alone. To assess performance in a downstream biomechanical context, we directly compared automated segmentations with those produced by three expert human annotators and constructed finite element models from each. Quantitative comparisons of segmentation accuracy and statistical analyses of finite element outputs demonstrate that automated segmentations perform within the range of expert human annotations, with no systematic bias in mechanical predictions. By explicitly linking geometric accuracy to biomechanical outcomes, this work establishes an end-to-end pipeline for validating automated segmentation in computational biology. The framework will be applicable to a wide range of musculoskeletal and palaeobiological contexts, including comparative anatomy, ageing and disease studies, and fossil or incomplete specimens, where scalable and mechanically faithful segmentation is essential.

**Author summary:** Many studies now use CT scanning to build computer models of bones and test how they perform under load. However, these analyses usually start with manual segmentation, where an expert traces the bone in thousands of image slices to create a 3D surface. This step is time-consuming and introduces uncertainty because experts do not always agree exactly on where boundaries lie. If segmentation changes the model shape, it may also change predicted biomechanics.

In this work, we test whether deep learning can automate segmentation while preserving biomechanical outcomes. We train an attention augmented 3D U-Net to segment the adult zebrafish lower jaw and compare its outputs directly to three expert manual segmentations. We assess similarity using complementary measures that reflect both overall overlap and boundary differences. We then build finite element models from each segmentation and compare predicted stress, strain, and displacement distributions.

We show that automated segmentations are as consistent as human experts and generate indistinguishable biomechanical predictions, without systematic bias. This supports scalable, repeatable pipelines for linking anatomy to function, and provides a general approach for validating automated segmentation in studies where downstream modelling is the goal.

## Introduction

Isolation (segmentation) of areas or volumes of interest from high-resolution imaging data is a critical computational bottleneck in quantitative morphology, biomechanics, and development(1–3). Detailed, accurate segmentation is vital for quantifying morphological traits, assessing developmental abnormalities, and understanding the biomechanics of functional adaptations(4–6). These processes underpin our ability to analyse genotype-phenotype relationships, examine evolutionary and developmental datasets, and explore the structural foundations of musculoskeletal health(7,8). Craniofacial structures present unique segmentation challenges due to their intricate geometry, often small size, and proximity to other anatomical features. These issues are particularly pronounced when comparing similar phenotypes, such as wildtype and mutant specimens, which often exhibit subtle yet biologically significant differences(9,10). Early methods for image segmentation relied on threshold techniques, boundary detection-based methods, region growing or a combination of boundary and region criteria(11). Early practitioners of segmentation developed mathematical tools(1,12–14) for image segmentation, but as scan quality increased, manual thresholding approaches by humans with expert knowledge of anatomy superseded strictly bound algorithmic approaches. This transition reflected the difficulty of developing universally applicable algorithms, particularly in developmental contexts where ossification patterns vary dramatically throughout development.

Despite its advantages for user-control, manual segmentation remains a labour-intensive and time-consuming task, often requiring expert annotators to achieve consistent and accurate results(2). Complex geometry such as craniofacial structures typically demands high levels of user input, making manual methods impractical for large-scale studies or high-throughput workflows. Additionally, manual segmentation lacks any formal error model, complicating downstream uncertainty estimations. This limitation is particularly problematic in research areas that require the analysis of large specimen datasets. For instance, in studies of skeletal development or craniofacial dysmorphology, large sample sizes are often needed to account for individual variability and genetic diversity within a population(15–17). Manual segmentation for these studies may introduce inter- and intra-operator variance as an unquantified source of computational noise. Where segmented geometry is used for further study, this could lead to potential biases analyses such as morphometrics or biomechanical analysis.

A solution is to apply deep learning auto-segmentation. Deep learning replaces subjective human interpretation with deterministic inference, but only if validated and trained on appropriate data(18). In recent years, the use of automated deep learning approaches in life sciences has advanced image segmentation(19,20), from biomedical to palaeontological sciences(21–26). Unlike traditional algorithms that rely on global thresholds or gradient-based edges, deep learning models emulate the contextual inference of expert annotators, combining intensity values with morphological features at different length scales to distinguish features(27). Convolutional neural networks (CNNs), particularly U-Net architectures, have become popular for biomedical image analysis because of their ability to capture complex spatial features across multiple scales. As such, the adoption of deep learning in biomedical segmentation, particularly through CNNs and U-Net-based architectures, has provided a foundation for accurate, reproducible segmentation tasks. U-Net architectures, initially developed for medical image segmentation, have been widely adapted to handle diverse biological data types, including MRI, CT, and micro-CT scans(19,20).

Attention mechanisms, which focus the model’s learning on critical regions of interest, enhance segmentation accuracy for anatomically complex structures by selectively weighting relevant features. This approach is especially valuable in craniofacial studies where the structures of interest may vary in size, orientation, or spatial relationship with adjacent tissues. Therefore, integrating these attention-based models into craniofacial segmentation workflows has the potential to achieve consistent, high-precision outputs, even in cases where traditional methods may fail due to variability in morphology. An additional advantage of attention-based U-Nets is their ability to perform out-of-sample inference, meaning they can generalize to new, previously unseen data(28,29). For example, by carefully training models on both wildtypes and mutants of the same species provides a framework suitable for studying the effects of genetic variation on craniofacial morphology.

Several issues remain with current approaches in deep learning for automated segmentation, Best practice in reporting results of automated image segmentation exists(30–32), yet few studies take up best practice advice. Methods for evaluation of model performance on geometrically complex biological shapes such as jaws or cranial bones require further interrogation to determine if reported percentage scores of model accuracy are “good-enough”. This applies to comparison with human manual segmentation outcomes and the impact on auto-versus human segmentation for downstream uses (e.g. morphometrics and biomechanics). Deep learning methods result in inaccuracies, but we still understand little of how DL inaccuracy compares to user-dependent inaccuracy for image segmentation tasks(33). This is the case for intra-user and multiple, inter-user segmentation results.

Furthermore, to our knowledge, no study has investigated this intra-user error effect on downstream usage of data, including biomechanical studies. Biomechanics research could benefit from automated segmentation, as the data generated can be directly fed into computational biomechanical models, including finite element (FE) models to simulate stress, strain, and other functional parameters(34–36). This could expand our capacity to investigate the functional morphology of craniofacial structures, linking geometric variation with function in a biologically meaningful way. Establishing a quantitative expert human benchmark is essential for evaluating and contextualising the performance of automated segmentation. In medical imaging, it is well recognised that inter- and intra-operator variability among expert annotators can be substantial, often exceeding the differences observed between expert and algorithmic outputs(37–39). These studies demonstrate that segmentation “error” is not a fixed property of an algorithm but a distribution that must be interpreted relative to human precision. In biological imaging, such reference frameworks are rarely established, leaving the field without clear criteria for determining when automated outputs are “good enough” for downstream analyses.

Here we address the problem of validating automated craniofacial segmentation in the absence of an absolute geometric ground truth yet downstream analyses that are sensitive to geometric variation. Using an attention-augmented 3D U-Net, we implement an automated, reproducible pipeline for segmenting the adult zebrafish mandible, which we use as a controlled benchmark system for testing segmentation fidelity and functional equivalence. We quantify segmentation uncertainty by comparing automated outputs to intra- and inter-operator variability among expert human annotators and evaluate geometric agreement using both volumetric (*Dice*) and surface-based (*Hausdorff*) metrics, which we integrate into a single conformity measure (the *Dice-Hausdorff* score). To assess whether geometric differences translate into differing functional interpretations, we propagate both manual and automated segmentations into finite element models and compare their biomechanical outputs. Zebrafish provide an experimentally tractable vertebrate system for this analysis due to their genetic accessibility, established use in musculoskeletal research(4,40–42), and suitability for high-resolution micro-CT imaging; however, finite element models of adult zebrafish jaws(43–45) have not previously been validated in this context. By quantifying the range of geometric and functional variability among expert human segmentations, we can define a biologically meaningful tolerance envelope, one that captures the natural uncertainty inherent to manual interpretation. Automated methods that perform within this envelope can thus be considered functionally equivalent to human performance, allowing validation to move beyond abstract geometric metrics toward biologically relevant, reproducible standards. Accordingly, this study aims to (i) quantify segmentation uncertainty relative to expert human variance, (ii) propose and evaluate a combined Dice–Hausdorff conformity metric for complex biological shapes, and (iii) test whether automated segmentation preserves downstream biomechanical predictions.

## Materials and methods

### Specimens

Zebrafish were raised at the University of Bristol, all animal work was approved locally by University of Bristol AWERB (Animal welfare and ethical review body) and performed under UK Home Office Project licence PP4700996. g

Zebrafish used ranged in age from 3 months to 3 years. Three months was chosen as the lower limit as this is the earliest stage we see substantial ossification (that is easily detectable via low resolution CT) throughout the whole jaw. Prior to this, mineral density is variable across the ossified elements. For the machine learning training dataset we selected 47 specimens, 6 at 3-6 months; 12 at 7-12 months; 10 at 13-18 months; 6 at 19-24 months; 6 at 25-30 months; 7 at 31 to 36 months (Supplementary Figure 1). For the comparative manual segmentation we used one 3 year old fish, towards the upper limit of our age range. This was to test the model against an ‘aged’ fish, where we anticipate less homogenous bone mineral density and a greater degree of modelling and remodelling to have occurred throughout the anterior of the head. All selected scans were screened for scan quality (lack of ring artifacts or very poorly resolved contrast).

### Scanning and pre-processing

Adult fish were fixed in 4% PFA for 1 week followed by sequential dehydration in 70% ethanol. Whole fish were batch scanned 10 fish per scan, using an XT H 225 ST micro-CT scanner (Nikon) with 21 µm voxel size using an X-ray source of 130□kV, 53□µA without additional filters. Gray values were calibrated with phantoms of known densities (0.25 and 0.75□g□cm3 of CaHA). After reconstruction of the tomographic images, 1999 slices were imported into ImageJ software(46,47) and converted from 16-bit greyscale to 32-bit greyscale tiff images to meet the requirements of our downstream processing.

### Deep Learning

#### Training Data

For efficient training, the CT image tiff stacks were manually cropped to a cube of 192×192×192 voxels around the region of interest. The network was trained on randomly extracted 160×160×160 voxel patches, a patch based approach waws selected so we could fine tune the model for future applications of other ROIs. Data augmentation (random rotations by multiples of 90°, reflections and affine transforms) was applied using TorchIO(48,49). The model was trained on 47 manually segmented datasets, segmented by one individual in Avizo 3D 2021.2(50) creating a labelled tiff stack of 192 slices around the jaw ROI. This dataset of 47 tiff stacks was composed of 32 full-jaw segmentations, and a supplementary dataset of 15 tiff stacks where only the jaw joint was labelled. This was because the region around the joint is the most difficult region to label, since zebrafish have a very complex skull morphology with many bones that have similar intensity values. The additional training data allows the model to learn the shape of the jaw joint without requiring the time-consuming annotation of 15 complete jaws. The jaw-label fields from the jaw segmentations were overlain onto the original 192×192×192 cropped CT tiff stacks in Python to create jaw label-CT slice masks. These masks were used as the deep learning training dataset.

### Segmentation Model

We used a 3D U-Net convolutional neural network (CNN) architecture with attention gates, implemented in PyTorch via MONAI(51), to segment the zebrafish jaw from micro-CT data. The U-Net is a fully convolutional neural network consisting of an encoding path and a decoding path, connected by attention-gated skip connections. The encoding path applies repeated convolution, activation, and pooling operations to down sample the input image and extract hierarchical features. The decoding path progressively up samples these features and concatenates them with attention-modulated feature maps from the encoding path. The attention mechanism in each skip connection selectively emphasizes features that are most relevant to the segmentation task while suppressing less informative activations(52). (Figure 2).

**Figure 1.**
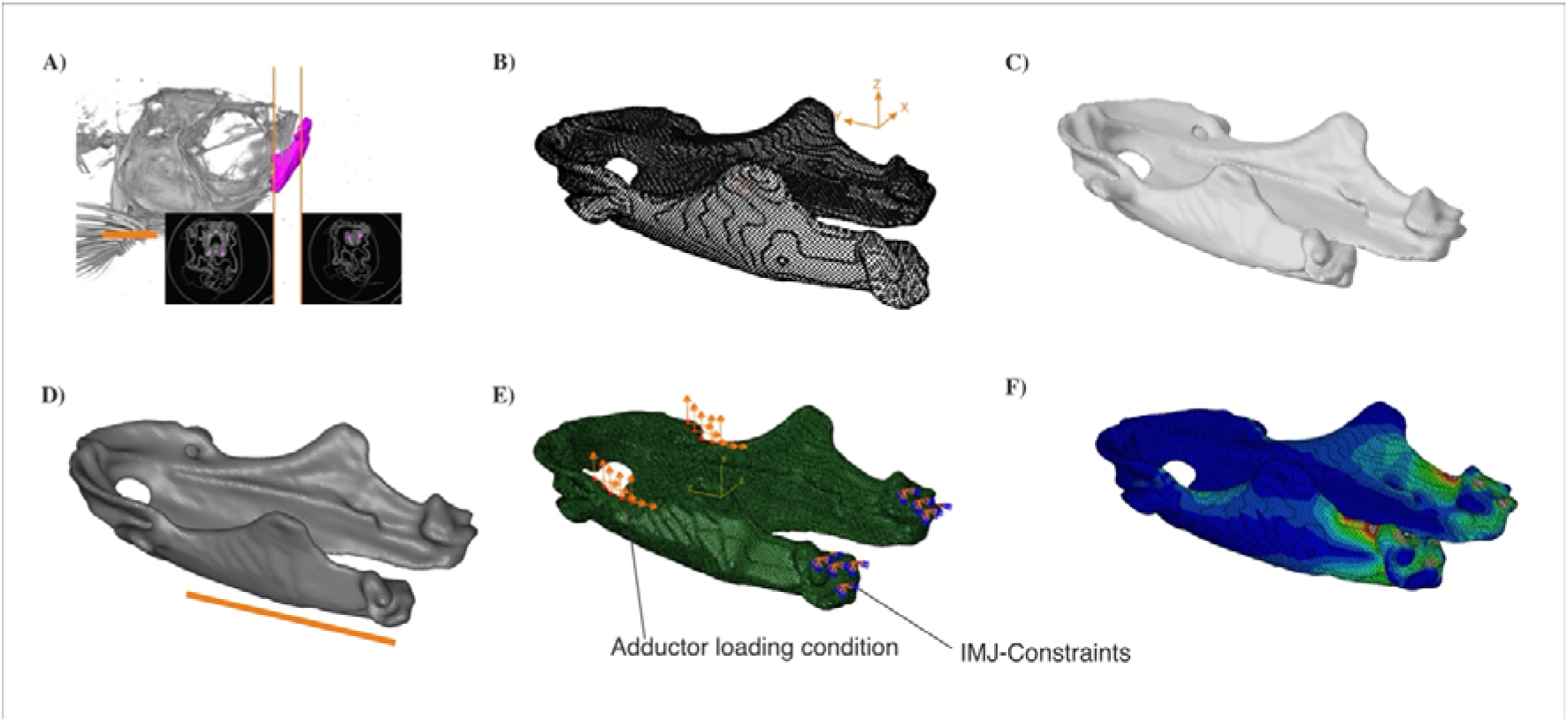
End-to-end workflow for finite element analysis (FEA) of the zebrafish lower jaw. Orange scale is 2mm. (A) Segmentation of the lower jaw is completed manually in Avizo or automatically via inference, followed by orientation in Blender using an anatomical coordinate system. Lower jaw target is highlighted in pink with representative tiff slices labelled below. (B) Models are checked for 3D print compatibility using Blender’s 3D Print Toolkit. (C) Mesh repair and optimization are completed in Geomagic Wrap using MeshDoctor. (D) Tetrahedral meshing is performed in Hypermesh, filling in the internal structures and converting the shell to a volume. (E) Boundary conditions such as constraining movement of the Inter-Mandibular Joint (IMJ) and material properties are assigned in Abaqus note adductor loading conditions are taken from Figure 2. (F) Resulting deformation and stress fields are visualized following simulation in Abaqus.

**Figure 2.**
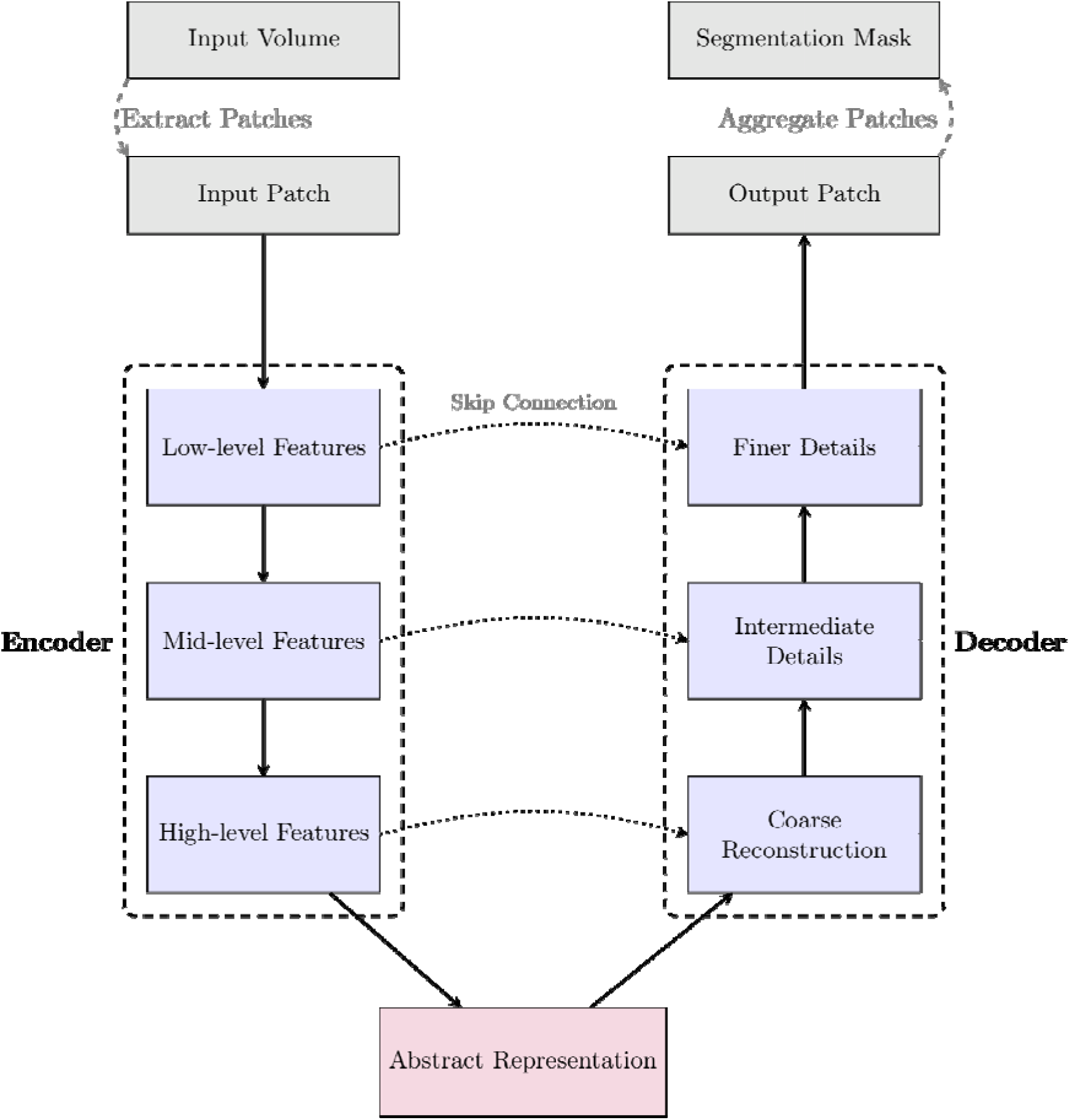
a schematic diagram of the U-Net architecture. Patches are extracted from the CT scan volume, which are then passed through an encoder-decoder network. The encoder extracts features at various scales, compressing information into a compact bottleneck representation. The decoder progressively reconstructs spatial details from coarse structures to fine details. Skip connections preserve spatial context by passing information directly from the encoder to the decoder. The output patches are aggregated to form the final segmentation mask.

The model was trained using the Adam optimiser with Pytorch’s default parameters on an NVIDIA RTX A6000 GPU. To address data imbalance, we used the Tversky loss function, which generalises the Dice coefficient by weighting false positives and false negatives. Jaw voxels comprised ∼0.1% of the total scan volume; the values α=0.2, β=0.8 were used in the loss to penalise false negatives so that the small structure of the jaw would be detected. To assess the uncertainty inherent in the model’s training process, we repeated the training process 20 times using the same training data, selecting the highest performing model from this process to present.

#### Surface and volume evaluation metrics

To assess the accuracy of the automated segmentations, we combined two common validation metrics: Dice score and Hausdorff distance.

The **Dice score** is a widely used measure of how much two volumes overlap. It gives a value between 0 and 1, where 1 indicates perfect agreement with the manual segmentation. While it is useful for summarising overall similarity, it can be misleading in cases where two shapes have similar volumes but different surface geometry (see Figure 3). This is especially common in thin or tapering structures like the zebrafish jaw.

**Figure 3.**
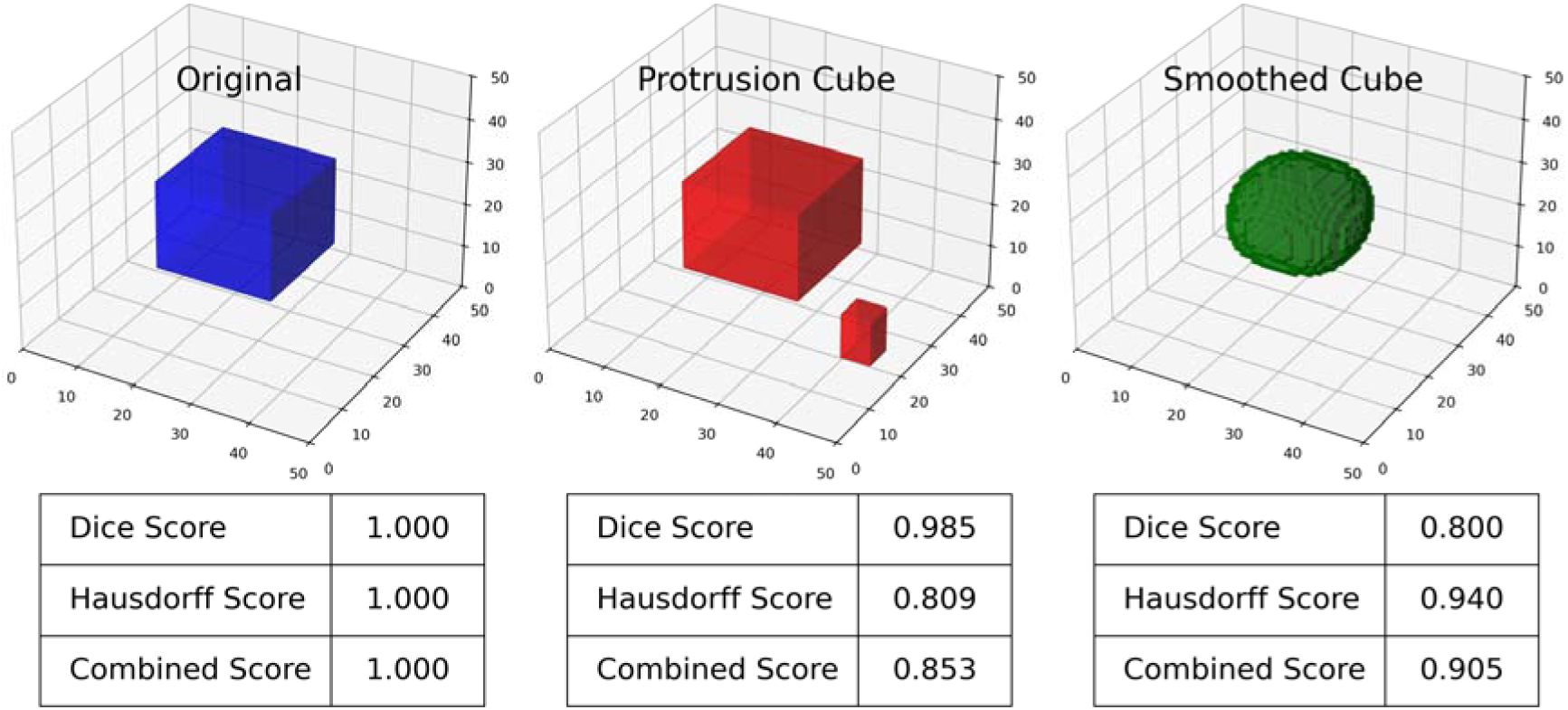
Comparison of three objects illustrating the utility of a combined score. Each column shows a 3D rendering with Dice similarity, Hausdorff score, and combined score relative to the original cube. The protrusion cube demonstrates the Dice score’s insensitivity to spatial disparity. The smoothed variant retains a relatively high Hausdorff score despite a large volumetric alteration, since the changes introduced by smoothing lie near the cube’s surface. The combined score is dependent on a scaling factor, but combines both volume and surface metrics.

The **Hausdorff distance** captures how far apart the surfaces of two shapes are. It identifies the maximum possible shortest distance between two objects, making it sensitive to mismatches in boundary shape and location. This is particularly important for applications like finite element analysis, where accurate surface geometry directly affects results (see Figure 3). The Hausdorff distance is a length which ranges from 0 to the maximum length in the image (i.e., the space diagonal of the volume). For ease of interpretation, here we introduce the Hausdorff score, defined as the Hausdorff distance scaled by its maximum value, transformed to take values between 0 and 1.

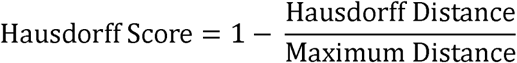

Because these each capture different aspects of segmentation quality—volume versus boundary—we introduce a **combined Dice-Hausdorff score**. This metric balances internal agreement with external shape accuracy. While not yet common in biological shape analysis, such combined metrics are recognised as important in high-precision workflows. In our context, it verifies that segmentations are not just broadly similar in size, but also closely matched in shape—both critical for biomechanical modelling. In practice, the Hausdorff score will be close to 1 for reasonable segmentations, so we use a scaling factor α to balance the contributions from the Dice and Hausdorff metrics. A value of 0.25 was used in this work.

Combined Score = α·Dice Score + (1-α)·Hausdorff Score

An illustration of the relative strengths and weaknesses of these metrics, see Figure 3, shows an example of modifications (addition of protrusions, smoothing) to a cube and the resulting effects on the Dice and Hausdorff metrics.

To assess the spread as well as the performance, we report the median and (2.5, 97.5)% quantiles for these metrics across the 20 trained models.

### 2.4. User Segmentation

We compared the results of the deep learning model to jaw shapes segmented by three human annotators. A jawbone CT scan that was not a part of the model’s validation set was segmented manually by three different annotators experienced in CT segmentation and craniofacial anatomy. One annotator (P1) had segmented the training dataset; another (P2) was familiar with fish anatomy; the third (P3) was familiar with tetrapod but not fish craniofacial anatomy. Scans were obtained from the same uCT scanner and using the same scan settings. Segmentation was performed in Avizo (2020.3).

### Finite Element Analysis

#### Finite Element Models

Finite element simulations were used to assess the biomechanical consequences of model shape variation due to jaw segmentation process. We tested how closely the segmentations—manual and automated—reproduce mechanical performance under loading conditions mimicking buccal feeding.

In Avizo, all segmented models were subjected to ‘Unconstrained smoothing’ applied to the Exterior, with a ‘Smoothing extent’ of 3.5. Segmented jaw surface meshes were exported from Avizo as .stl files and imported into Blender (blender 4.4), where they were aligned to a standard Cartesian coordinate system. Then resampled to a uniform mesh resolution using triangular surface faces with an edge length of 0.03 mm². The 3D Print Toolkit was used to identify and correct common mesh issues, including non-manifold edges, sharp faces, intersecting geometry, and incorrect normal orientation. Surface meshes were then exported from Blender and further refined in Geomagic Wrap (v2024.3.1) using the Mesh-Doctor tool. Next, surface meshes were imported to Hypermesh (v.2023.1 Altair), where they were converted to volumetric meshes made up of quadratic tetrahedral elements (C3D10). For finite element analysis, a mesh must be watertight, manifold, free of intersecting or inverted elements, and composed of high-quality tetrahedral elements so the quality of the final volumetric mesh, including maximum aspect ratio and Jacobian metrics, was checked.

Vertex, edge, face, and triangle counts were checked for consistency across all models, with mesh statistics presented in Table 1.

**Table 1.** Mesh model. Summary of complexity by mesh topology metrics for each segmentation model, including the number of vertices, edges, faces, and triangles used in finite element analysis. The Inference model is derived from automated segmentation, while P1, P2, and P3 represent manually segmented human models.

| Annotator | Vertices | Edges | Faces |
| --- | --- | --- | --- |
| P1 | 32194 | 64396 | 32198 |
| P2 | 31870 | 63748 | 31874 |
| P3 | 33904 | 67816 | 33908 |
| Inference | 32658 | 65328 | 32664 |

#### Loading

Each finite element model was assigned isotropic, homogeneous material properties, with a Young’s modulus of 7 GPa and a Poisson’s ratio of 0.3. These values reflect an approximation of the composite nature of zebrafish bone(53) and were chosen to ensure mechanical behaviour consistent with physiological estimates from Hammond Lab experiments (Supplementary Materials).

In Abaqus, loading and boundary conditions were applied to simulate buccal feeding. The model was constrained at the intramandibular joints (IMJs) with five surface nodes placed on either side that were constrained in all translational and rotational degrees of freedom (U1-U3, UR1-UR3). Loading conditions replicating the bilateral contraction of the adductor mandibulae muscles were applied at five insertion points along each side of the jaw (see Figure 1D). A total of 0.5 N was applied symmetrically at five insertion points per side, with each node receiving a 0.05 N vector force at 45° to both the X and Y axes, in line with physiological estimates from Hammond Lab experiments (Supplementary Materials). Static linear analyses were run using the *Static, General* step in Abaqus. Nonlinear geometric effects were assumed negligible.

#### Analysis and comparison metrics

FE outputs were assessed quantitatively by visualising von Mises stress (MPa) magnitudes and distributions using a uniform scale of banded colours. Outputs were also assessed quantitatively via nodal deformation, mean von Mises stress (MPa), and maximum strain (*E*). Similarity was assessed between human and automatically segmented models using multiple statistical approaches. The Kolmogorov-Smirnov (KS) test was applied to compare strain and stress distributions. As a non-parametric test, KS is particularly suitable for assessing differences in the shape of cumulative distributions without assuming normality or requiring identical sample sizes. This allowed us to test whether the observed differences in strain or stress patterns reflected meaningful biomechanical variation or were within the bounds of distributional similarity. For displacement fields, which often vary in node count and lack pointwise correspondence between segmentations, we used the first Wasserstein distance (also known as Earth Mover’s Distance). The Wasserstein distance accounts for both the magnitude and spread of displacement values, offering an interpretable measure of the “effort” required to transform one distribution into another. This makes it well-suited for comparing anatomically matched but topologically distinct meshes. The method has seen increasing use in biomedical shape and distributional analysis where geometric inconsistency and sampling variation are present(54,55). As all segmentations derived from the same individual specimen and mesh resolution was standardized, formal mesh convergence testing was not required.

## Results

### Segmentation

Label fields from the three human annotators and the inference from the U-Net model can be seen in Figure 4. A representative slice through the scan is shown, depicting the jaw joint.

**Figure 4.**
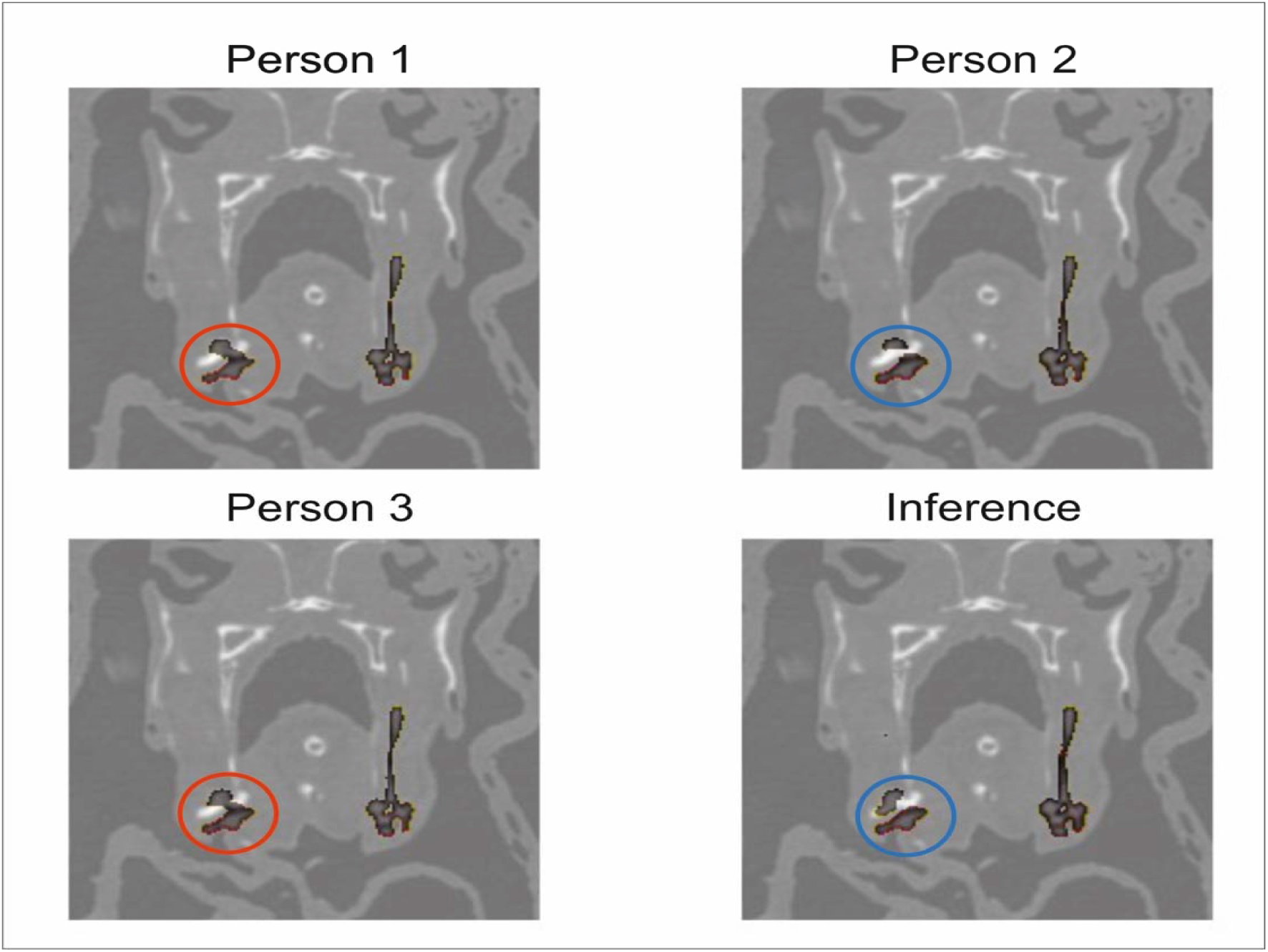
a representative slice through the CT scan. The segmentation mask (black) for this slice is overlaid on the CT scan (greyscale). All four segmentations are similar; differences appear near the left jaw joint (circled). P1 and P3 show a contiguous segmentation (red, solid circle); P2 and the model show separated regions in this slice (blue, dashed circle). The right side of the segmentation mask appears larger because of misalignment between the fish and the CT beamline, due to the way in which fish are loaded for scanning.

**Figure 5.**
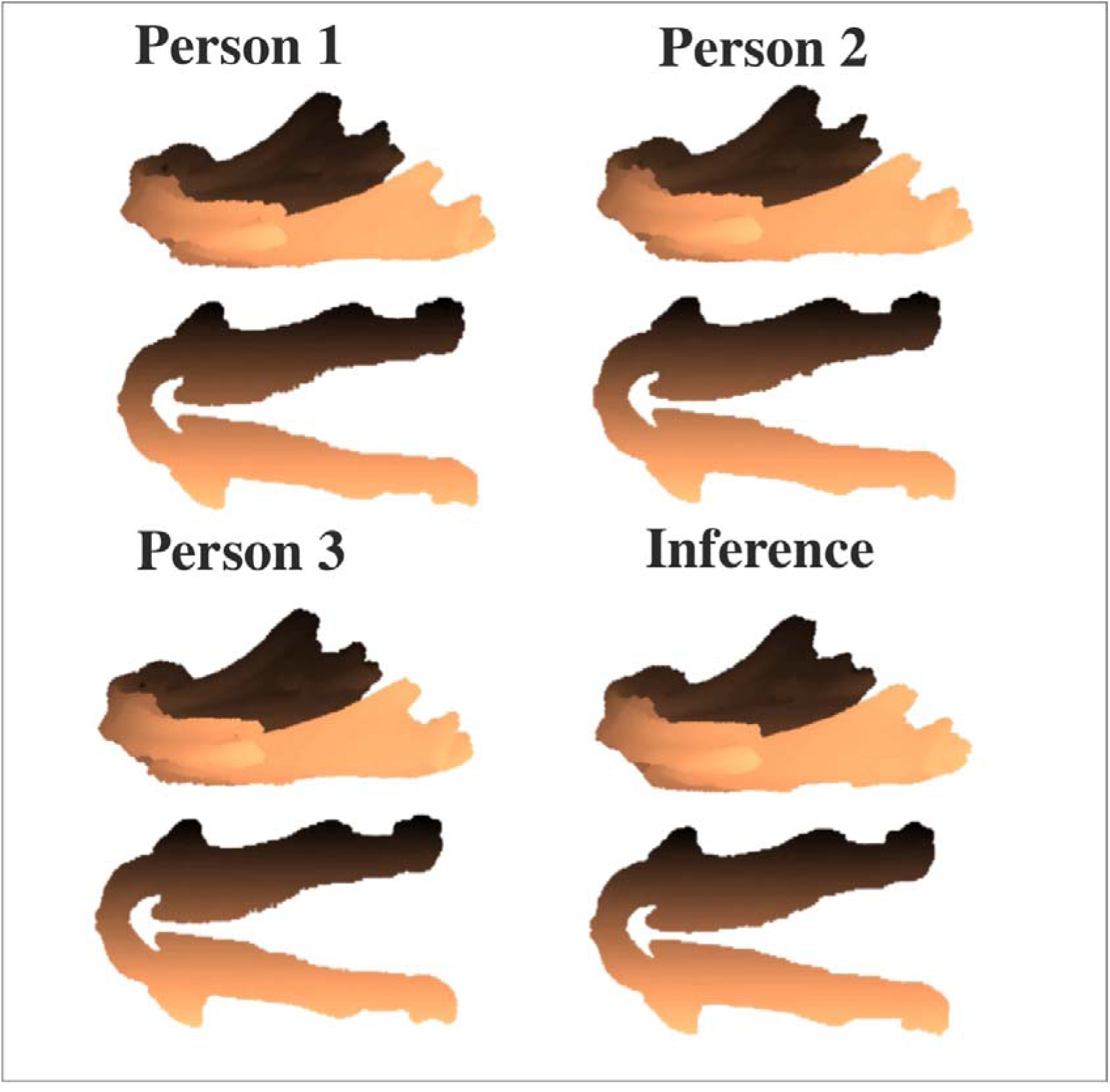
projections of the segmentations performed by the human annotators and the model. Skewed lateral view presented above and ventral beneath for each result. Note the base of the jaw appears to have the greatest variance where there is greatest contact and overlap with other bone material. The anterior of each of these has exceptionally consistent results as thresholding could prove more effective.

**Figure 6.**
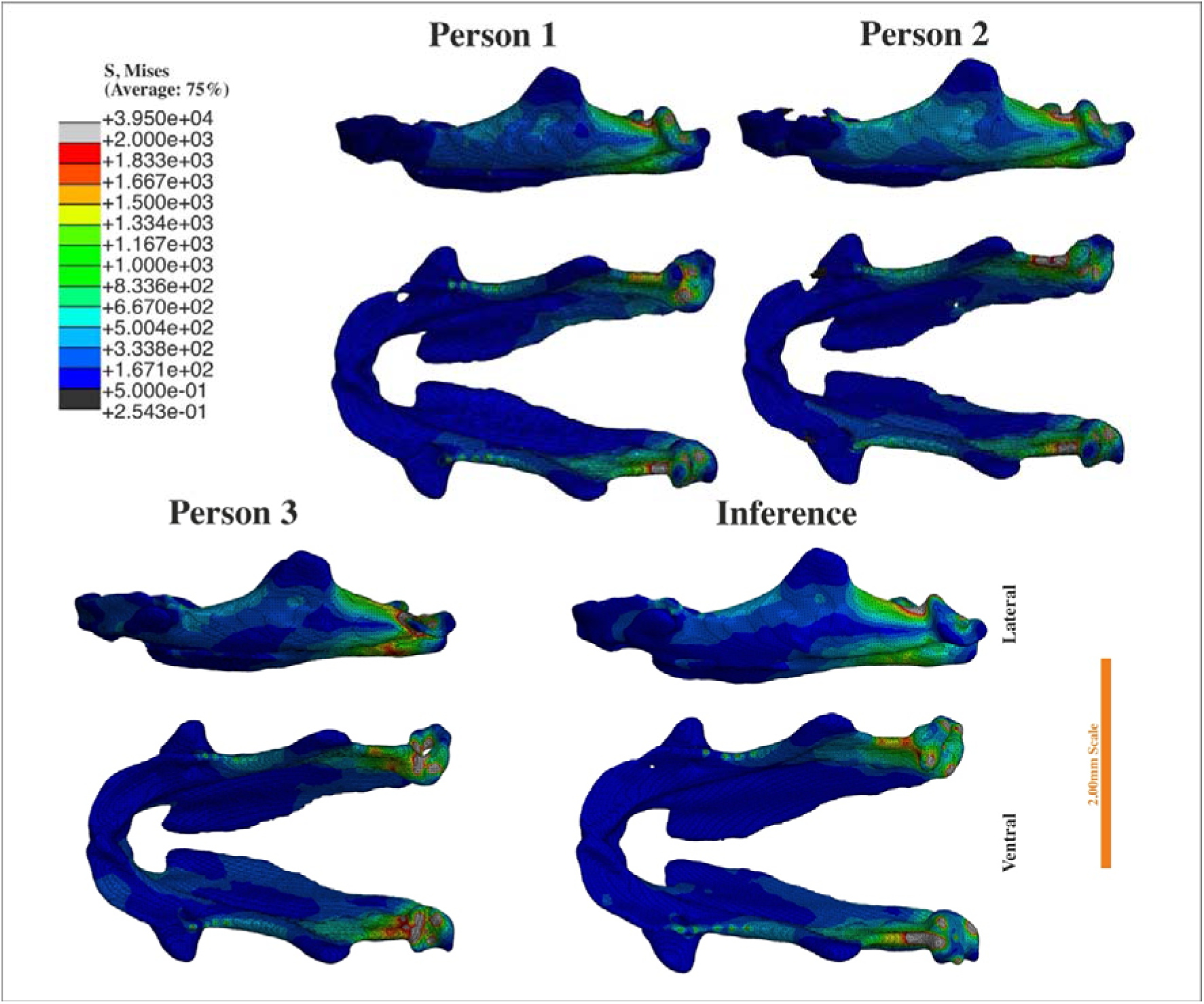
Finite element von Mises stress distribution across four manual segmentations and one automated segmentation. Colour maps show stress distributions (MPa) under identical loading conditions for three human segmentations (Person 1-3) and the automated inference model. Despite minor geometric differences, all segmentations yield comparable stress patterns, particularly in the anterior. Scale bar-2.00 mm. Lateral view presented above ventral below.

This is a difficult region to segment, since several bones are present, butting on to one another.

### Segmentation and finite element analysis metrics

The metrics from the segmentation and Finite Element Analysis tests can be seen in Table 2. The histograms from which the finite element metrics were derived can be seen in Supplementary Figure 4.

**Table 2.** Performance metrics for the segmentation and FE. P1 segmented the same jaw three times; (the third value in the table being the ‘ground truth’) and the resulting FE was used as the ground truth. The median and 2.5%/97.5% quantiles of the segmentation metrics are reported, after training 20 models on the same training data. Only one auto-segmentation was taken through to FE, the parameters from which are reported in the table. The intra-human and inter-model variation is similar, and both distributions overlap significantly in all segmentation metrics.

|  | *Segmentation Metrics* |  |  | *FE Metrics* |  |  |
| --- | --- | --- | --- | --- | --- | --- |
|  | *Dice* | *Hausdorff* | *Dice-Hausdorff* | *Displacement*<br>*Wasserstein*<br>*Distance* | *Strain*<br>*KS* | *Stress KS* |
| P1 (3 trials) | 0.90, 0.93, 1.0 | 0.98, 0.98, 1.0 | 0.96, 0.97, 1.0 | 0.00 | 0.00 | 0.00 |
| P2 | 0.98 | 0.99 | 0.99 | 0.18 | 0.10 | 0.12 |
| P3 | 0.95 | 0.98 | 0.97 | 0.32 | 0.15 | 0.13 |
| Inference<br>(median<br>[IQR]) | 0.91 (0.89-0.95) | 0.98 (0.98-0.98) | 0.96 (0.95-0.97) | 0.20 | 0.02 | 0.06 |

Quantitative shape comparison using Dice and Hausdorff-derived metrics demonstrate the automated segmentation performs within the range of human segmentations. The Dice coefficient and 1-Hausdorff scores show that all models, including the automated one, are closely aligned, with particularly high overlap between P1 and P2. The automated model achieved Dice, Hausdorff, and Dice–Hausdorff scores of 0.98, 0.99, and 0.99 respectively, outperforming the intra-operator variation observed in Person 1’s repeated segmentations.

This represents a remarkable result, given that Person 1 also produced the training dataset, indicating that the model generalises beyond the variability of its own ground-truth source

### 3.3 Biomechanical comparisons

Qualitatively, all models show comparable stress distributions with higher stresses observed at the intramandibular joint, posterior of the coronoid process (dorsally and ventrally), and at modelled muscle attachment sights. However, there are small differences. The Person 2 model shows relatively higher stress between the two processes, likely a result of reduced geometry due user thresholding selection during segmentation. Additionally, there are small variations in stress distribution at the intramandibular joints, which is likely due to user knowledge of this complex anatomy.

Quantitative comparisons of stress and strain distributions using the Kolmogorov-Smirnov test showed that the automated segmentation produced outputs within the range of reference (P1) and the other expert annotators (Table 2). For both strain and stress, the lowest KS statistic was observed for the automated model indicating strong biomechanical convergence (Table 2).

Despite slight geometric variation, particularly in bulkiness around the margins, the automated segmentation preserved critical features influencing load transmission. Displacement field comparisons using Wasserstein distance further support this, with the inference model showing lower displacement divergence from Person 1 than one of the human annotator derived models (Table 2). This suggests that minor morphological deviations introduced by the automated pipeline do not substantially affect the biomechanical interpretation.

## Discussion

### Material properties and model assumptions

Finite element analysis provides a quantitative means of testing whether automated segmentations preserve biologically meaningful geometry and functional inference(56,57). To isolate segmentation performance from material complexity, all models were standardised using isotropic, homogenous bone properties (E = 7 GPa; *v* = 0.35). these parameters lie within the empirically measured range for acellular teleost bone and are consistent with other teleost analyses. They approximate the composite stiffness of zebrafish bone, which is less mineralised than mammalian cortical tissue. The elevated Poisson’s ratio reflects the increased ductility of small teleost bones.

### Biomechanical patterns and biological interpretation

Predicted strain and stress distributions in the zebrafish lower jaw were consistent with expected loading regimes during mouth opening and closure. Peak von Mises stresses occurred around the retro-articular process and posterior joints, with elevated strain along the dorsal Meckel’s cartilage and at muscular insertion sites. These areas correspond to known attachments of the intermandibular is musculature and loading, indicating that the finite-element models reconstructed biologically realistic load paths(42,58). Comparisons among human-segmented and model-derived meshes revealed that the principal axes of deformation were conserved across all datasets. Despite minor geometric variation between operators and automated outputs, the orientation and distribution of strain remained stable, demonstrating that the overall functional topology of loading is robust to boundary definition. This consistency provides direct evidence that automated segmentation preserves biomechanical fidelity.

### Methodological integration: segmentation to FEA

Automated segmentation and biomechanical modelling are rarely combined in vertebrate research. automated segmentation models have been successfully trained to work on We believe that validation is best completed on at least a representative downstream analysis intended of the pipeline. The workflow developed here bridges that gap by coupling an attention-augmented 3D U-Net with automated finite-element generation. This approach converts imaging data directly into functional models, standardising each stage of analysis and enabling objective comparison across specimens, genotypes, and developmental stages.

Validation suggests that models can be compared to a ground truth, but segmentations are by nature and at best an expert’s interpretation of contrast-bound structures in a μCT stack(2). Segmentation is applied in many cases to computational experiments such as finite element analysis with strict controls on many downstream processes such as meshing, but with relatively little regard for the repeatability in the human inference stage. With only a sole practitioner determining the final output, there is no way to assess the precision of both the morphology or functional output. By having multiple experts segment the same structure, we can generate a space that has similar functional outputs and can assess the model that we generate to these results.

The inclusion of Hausdorff with the Dice score summarises the variability of the surface shape consistency with the overlap per slice well. The results of this are corroborated directly by the assessment of functional performance as both times, the U-Net annotations were within the tolerance envelope established by human annotators. One major benefit of model segmentation is the labels generated by the U net are deterministic, and the result is entirely repeatable, along as it is not retrained, even on the same training data, the resultant labels will be the same every time it is implemented on the same tiff stack. The training process itself is not deterministic, and following retraining, even on the same training set will result in a differently performing model.

Inter- and intra-human comparisons demonstrated that segmentation variability among experts, and between experts and the trained network, was low and did not substantially affect downstream mechanical results. Minor boundary differences in low-contrast or geometrically complex regions produced negligible changes in stress or strain orientation. The U-Net, trained on expert-labelled exemplars, captured the mean interpretive behaviour of human annotators and produced deterministic outputs within the natural range of intra-human variability. Because no absolute ground truth exists for biological segmentation, precision is a more meaningful performance criterion than nominal accuracy. The FEA results from the automated segmentation yield mechanical outputs statistically and functionally equivalent to those derived from manual meshes.

This convergence establishes FEA as an independent validation test of segmentation fidelity. If automated segmentations generate strain fields indistinguishable from human-derived models, they can be considered functionally accurate even without a definitive geometric reference. Hausdorff and Dice metrics provide complementary geometric measures—surface deviation and volumetric overlap—but FEA integrates these assessments within a biologically relevant, functional framework. Additional variation may arise from meshing or landmark placement, suggesting that future pipelines should incorporate automated or learned constraint detection to further stabilise the analysis process.

By integrating segmentation metrics with mechanical outcomes, this study provides the first fully validated segmentation-to-FEA pipeline for zebrafish jaws. The resulting functional measures—mean strain energy density, stiffness, and deformation asymmetry—represent quantitative phenotypes that can be compared across individuals or experimental groups.

Because segmentation and meshing are fully algorithmic, identical parameters can be applied across datasets, facilitating reproducible analyses across developmental, genetic, and environmental contexts.

### Comparison to other auto-segmentation studies in medical, biological, and palaeontological fields

Few studies have investigated variation between human practitioners of segmentation, despite its critical influence on downstream quantitative analyses. In medical imaging, inter-operator variability has been shown to affect diagnostic and prognostic metrics, motivating the widespread use of automated methods for tumour delineation, organ segmentation, and bone reconstruction(59,60). However, even in clinical contexts, validation typically focuses on geometric concordance—Dice, Jaccard, or Hausdorff distance—rather than assessing how segmentation differences propagate into functional models or biomechanical predictions. To address this imbalance, loss functions such as the Tversky loss have been developed to weight false positives and false negatives asymmetrically, improving model sensitivity to boundary precision in anatomically complex regions(61). Our presented work extends these frameworks by integrating both geometric and mechanical validation within a single analytical chain, establishing finite element outputs as biologically relevant performance benchmarks

In comparative and developmental biology, automated segmentation remains less established, often limited by smaller datasets and the complex, low-contrast morphology of μCT specimens(57,62–66). Previous studies in teleost, amphibians, and amniotes have applied deep learning to accelerate volumetric labelling, yet few have demonstrated equivalence to expert segmentations across both morphology and function(62,64,67). Outside of micropalaeontology, deep learning methods applied lack the versatility to work on multiple specimens, and at best accelerate the rate of segmentation that likely requires downstream cleanup pre meshing. Our results show that when trained on expert-annotated exemplars, attention-augmented architectures can generalise across fine-scale craniofacial structures while maintaining biomechanical fidelity. This supports the use of machine learning not merely as a convenience for throughput, but as a rigorous analytical tool that standardises human interpretation.

Palaeontological applications present an even greater challenge due to taphonomic alteration, matrix interference, and the absence of soft-tissue reference data. Emerging work using convolutional and transformer-based networks for fossil segmentation has achieved high geometric accuracy(65,66), yet functional validation remains largely unexplored. The framework presented here demonstrates that coupling segmentation with FEA provides an empirical means to assess whether reconstructed morphologies are mechanically plausible, offering a pathway for verifying automated reconstructions of fossil taxa where no living analogue exists. As palaeo-biomechanics increasingly incorporates digital reconstruction, automated segmentation validated by mechanical performance will become essential for ensuring that inferred behaviours and functional morphologies rest on reproducible geometric foundations.

### Limitations and future directions

The present models assume homogeneous, isotropic material properties and static loading conditions. In vivo, the zebrafish jaw experiences dynamic, multi-directional forces transmitted through muscle fibres, ligaments, and cartilage. Incorporating muscle wrapping, soft-tissue constraints, and time-dependent material behaviour would increase biological realism and enable dynamic validation tests under physiological conditions.

Although the segmentation pipeline produces reproducible meshes, geometric fidelity depends on CT resolution and training-set diversity. Multi-scale architectures combining 2.5D and 3D representations could improve delineation of thin or low-contrast structures such as the retro-articular process. Validation against digital histology or confocal reconstructions would further confirm boundary accuracy. Quantifying how segmentation uncertainty propagates into biomechanical variance remains a key challenge; ensemble analyses using multiple human and automated segmentations will help define tolerance thresholds for functional equivalence.

### Conclusions

This study presents the first automated segmentation and finite element (FE) analysis of the adult zebrafish jaw, providing a reproducible digital framework for quantifying craniofacial biomechanics in a key vertebrate model. By coupling attention-based deep learning with mechanical simulation, we demonstrate that automated segmentations preserve biologically meaningful geometry and generate stress and strain distributions that are statistically and functionally equivalent to those derived from expert human meshes. Finite element analysis therefore serves not only as a tool for biomechanical inference but as an independent, quantitative test of segmentation fidelity. Using both volumetric (Dice) and surface-based (Hausdorff) metrics, we illustrate the value of integrating complementary geometric measures with functional validation. The Tversky loss function combined with attention enables the model to achieve the results in the measured metrics despite the data imbalance in μCT scans. The uncertainty within the model’s predictions falls within the natural range of human variation, confirming that automated outputs are biomechanically indistinguishable from manual segmentations.

Beyond establishing the first validated digital models of the zebrafish jaw, this framework provides a scalable, transferable approach for evaluating and applying automated segmentation across vertebrate morphology. It enables high-throughput, functionally grounded phenotyping that links genotype, form, and biomechanical performance in evolutionary and developmental contexts.

## Acknowledgements

The work benefited from support provided by the Jean Golding Institute for Data Science and computational facilities within the Bristol Palaeobiology Research Group, as well as discussion with the Hammond and Rayfield lab groups.

## Code availability and Supplementary information

All code and supplementary materials are available through the following repository. https://github.com/JGIBristol/zebrafish_jaw_segmentation

## References

1. Pham DL, Xu C, Prince JL. Current Methods in Medical Image Segmentation1. Annual Review of Biomedical Engineering. 2000 Aug 1;2(Volume 2, 2000):315–37.

2. Zhang L, Tanno R, Xu MC, Jin C, Jacob J, Ciccarelli O, et al. Disentangling Human Error from the Ground Truth in Segmentation of Medical Images [Internet]. arXiv; 2020 [cited 2025 June 26]. Available from: http://arxiv.org/abs/2007.15963

3. Rayfield EJ. Finite Element Analysis and Understanding the Biomechanics and Evolution of Living and Fossil Organisms. Annual Review of Earth and Planetary Sciences. 2007 May 30;35(Volume 35, 2007):541–76.

4. Inbar Ben-Zvi, David Karasik, Cheryl Ackert-Bicknell. Zebrafish as a Model for Osteoporosis: Functional Validations of Genome-Wide Association Studies. 2023 Nov 16;

5. Graham J. Slater, Slater GJ, Blaire Van Valkenburgh, Van Valkenburgh B. Allometry and performance: the evolution of skull form and function in felids. Journal of Evolutionary Biology. 2009 Nov 1;22(11):2278–87.

6. Kague E, Karasik D. Functional Validation of Osteoporosis Genetic Findings Using Small Fish Models. Genes. 2022 Jan 30;13(2):279–279.

7. Schilling TF, Kimmel CB. Musculoskeletal patterning in the pharyngeal segments of the zebrafish embryo. Development. 1997 Aug;124(15):2945–60.

8. Fiedler I, Zeveleva S, Duarte A, Zhao X, Depalle B, Cardoso L, et al. Microstructure, Mineral and Mechanical Properties of Teleost Intermuscular Bones. J Biomech. 2019 Sept 20;94:59–66.

9. Hark R, Zürlein S, Nguyen VT, Gust G, Hekel L, Liedtke D. AI-assisted phenotyping in a zebrafish hypophosphatasia model enables early and precise detection of skeletal alterations. Sci Rep. 2025 Sept 17;15(1):32578.

10. Prendergast A, Ziganshin BA, Papanikolaou D, Zafar MA, Nicoli S, Mukherjee S, et al. Phenotyping Zebrafish Mutant Models to Assess Candidate Genes Associated with Aortic Aneurysm. Genes (Basel). 2022 Jan 10;13(1):123.

11. Csurka G, Volpi R, Chidlovskii B. Semantic Image Segmentation: Two Decades of Research [Internet]. arXiv; 2023 [cited 2025 Oct 21]. Available from: http://arxiv.org/abs/2302.06378

12. Adams R, Bischof L. Seeded region growing. IEEE Trans Pattern Anal Machine Intell. 1994 June;16(6):641–7.

13. Otsu N. A Threshold Selection Method from Gray-Level Histograms. IEEE Transactions on Systems, Man, and Cybernetics. 1979 Jan;9(1):62–6.

14. Canny J. A Computational Approach to Edge Detection. IEEE Transactions on Pattern Analysis and Machine Intelligence. 1986 Nov;PAMI-8(6):679–98.

15. McGurk PD, Lovely CB, Eberhart JK. Analyzing Craniofacial Morphogenesis in Zebrafish Using 4D Confocal Microscopy. J Vis Exp. 2014 Jan 30;(83):51190.

16. Schilling TF. Genetic analysis of craniofacial development in the vertebrate embryo. BioEssays. 1997;19(6):459–68.

17. Maili L, Ruiz OE, Kahan PH, Chiu F, Larson ST, Hashmi SS, et al. Facial analytics based on a coordinate extrapolation system (zFACE) for morphometric phenotyping of developing zebrafish. Dis Model Mech. 2023 June 1;16(6):dmm049868.

18. Neves CA, Tran ED, Kessler IM, Blevins NH. Fully automated preoperative segmentation of temporal bone structures from clinical CT scans. Sci Rep. 2021 Jan 8;11(1):116.

19. Olaf Ronneberger, Ronneberger O, Philipp Fischer, Fischer P, Thomas Brox, Brox T. U-Net: Convolutional Networks for Biomedical Image Segmentation. 2015 Oct 5;234–41.

20. Alex Krizhevsky, Krizhevsky A, Ilya Sutskever, Sutskever I, Geoffrey E. Hinton, Hinton GE. ImageNet classification with deep convolutional neural networks. Communications of The ACM. 2017 May 24;60(6):84–90.

21. Goswami A, Clavel J. Morphological evolution in a time of phenomics. Paleobiology. 2025 Feb;51(1):195–213.

22. Toulkeridou E, Gutierrez CE, Baum D, Doya K, Economo EP. Automated segmentation of insect anatomy from micro-CT images using deep learning. Natural Sciences. 2023;3(4):e20230010.

23. Lösel PD, van de Kamp T, Jayme A, Ershov A, Faragó T, Pichler O, et al. Introducing Biomedisa as an open-source online platform for biomedical image segmentation. Nat Commun. 2020 Nov 4;11(1):5577.

24. Mulqueeney JM, Searle-Barnes A, Brombacher A, Sweeney M, Goswami A, Ezard THG. How many specimens make a sufficient training set for automated three-dimensional feature extraction? Royal Society Open Science. 2024 June 19;11(6):rsos.240113.

25. Yu C, Qin F, Watanabe A, Yao W, Li Y, Qin Z, et al. Artificial intelligence in paleontology. Earth-Science Reviews. 2024 May 1;252:104765.

26. Didziokas M, Pauws E, Kölby L, Khonsari RH, Moazen M. BounTI (boundary-preserving threshold iteration): A user-friendly tool for automatic hard tissue segmentation. Journal of Anatomy. 2024;245(6):829–41.

27. Deep Learning for Semantic Segmentation | SpringerLink [Internet]. [cited 2025 July 15]. Available from: https://link.springer.com/chapter/10.1007/978-3-030-74478-6_3?utm_source=chatgpt.com

28. Xie Y, Yang B, Guan Q, Zhang J, Wu Q, Xia Y. Attention Mechanisms in Medical Image Segmentation: A Survey [Internet]. arXiv; 2023 [cited 2025 July 15]. Available from: http://arxiv.org/abs/2305.17937

29. Zhang J, Chen X, Yang B, Guan Q, Chen Q, Chen J, et al. Advances in attention mechanisms for medical image segmentation. Computer Science Review. 2025 May 1;56:100721.

30. Why rankings of biomedical image analysis competitions should be interpreted with care | Nature Communications [Internet]. [cited 2025 July 15]. Available from: https://www.nature.com/articles/s41467-018-07619-7

31. Luo W, Phung D, Tran T, Gupta S, Rana S, Karmakar C, et al. Guidelines for Developing and Reporting Machine Learning Predictive Models in Biomedical Research: A Multidisciplinary View. Journal of Medical Internet Research. 2016 Dec 16;18(12):e5870.

32. Seghier ML. Ten simple rules for reporting machine learning methods implementation and evaluation on biomedical data. International Journal of Imaging Systems and Technology. 2022;32(1):5–11.

33. Lallensack JN, Romilio A, Falkingham PL. A machine learning approach for the discrimination of theropod and ornithischian dinosaur tracks. Journal of The Royal Society Interface. 2022 Nov 9;19(196):20220588.

34. Bissinger O, Götz C, Wolff KD, Hapfelmeier A, Prodinger PM, Tischer T. Fully automated segmentation of callus by micro-CT compared to biomechanics. J Orthop Surg Res. 2017 July 11;12(1):108.

35. Brunt LH, Roddy KA, Rayfield EJ, Hammond CL. Building Finite Element Models to Investigate Zebrafish Jaw Biomechanics. Journal of Visualized Experiments (JoVE). 2016 Dec 3;(118):e54811.

36. Ramos-Soto O, Jo HC, Zawadzki RJ, Kim DY, Balderas-Mata SE. Automated Segmentation and Morphometry of Zebrafish Anterior Chamber OCT Scans. Photonics. 2023 Sept;10(9):957.

37. Warfield SK, Zou KH, Wells WM. Simultaneous Truth and Performance Level Estimation (STAPLE): An Algorithm for the Validation of Image Segmentation. IEEE Trans Med Imaging. 2004 July;23(7):903–21.

38. Metrics for evaluating 3D medical image segmentation: analysis, selection, and tool | BMC Medical Imaging | Full Text [Internet]. [cited 2025 Oct 21]. Available from: https://bmcmedimaging.biomedcentral.com/articles/10.1186/s12880-015-0068-x

39. Reinke A, Tizabi MD, Sudre CH, Eisenmann M, Rädsch T, Baumgartner M, et al. Common Limitations of Image Processing Metrics: A Picture Story [Internet]. arXiv; 2023 [cited 2025 Oct 21]. Available from: http://arxiv.org/abs/2104.05642

40. Bergen DJM, Kague E, Hammond CL. Zebrafish as an emerging model for osteoporosis: a primary testing platform for screening new osteo-active compounds. Frontiers in Endocrinology. 2019 Jan 29;10:6–6.

41. Busse B, Galloway JL, Gray RS, Harris MP, Kwon RY. Zebrafish: An Emerging Model for Orthopaedic Research. J Orthop Res. 2020 May;38(5):925–36.

42. Josepha Godivier, Lawrence EA, Mengdi Wang, Hammond CL, Nowlan NC. Growth orientations, rather than heterogeneous growth rates, dominate jaw joint morphogenesis in the larval zebrafish. Journal of Anatomy. 2022 May 5;

43. Brunt LH, Norton JL, Bright JA, Rayfield EJ, Hammond CL. Finite element modelling predicts changes in joint shape and cell behaviour due to loss of muscle strain in jaw development. J Biomech. 2015 Sept 18;48(12):3112–22.

44. Ofer L, Dean MN, Zaslansky P, Kult S, Shwartz Y, Zaretsky J, et al. A novel nonosteocytic regulatory mechanism of bone modeling. PLoS Biol. 2019 Feb;17(2):e3000140.

45. Lawrence EA, Aggleton J, van Loon J, Godivier J, Harniman R, Pei J, et al. Exposure to hypergravity during zebrafish development alters cartilage material properties and strain distribution. Bone Joint Res. 2021 Feb 1;10(2):137–48.

46. ImageJ Wiki [Internet]. [cited 2025 July 15]. ImageJ. Available from: https://imagej.github.io/software/imagej/index

47. Schneider CA, Rasband WS, Eliceiri KW. NIH Image to ImageJ: 25 years of image analysis. Nat Methods. 2012 July;9(7):671–5.

48. Pérez-García F, Sparks R, Ourselin S. TorchIO: a Python library for efficient loading, preprocessing, augmentation and patch-based sampling of medical images in deep learning [Internet]. Computer Methods and Programs in Biomedicine. 2021 [cited 2025 Oct 28]. Available from: https://www.sciencedirect.com/science/article/pii/S0169260721003102

49. Pérez-García F, Sparks R, Ourselin S. TorchIO: A Python library for efficient loading, preprocessing, augmentation and patch-based sampling of medical images in deep learning. Computer Methods and Programs in Biomedicine. 2021 Sept 1;208:106236.

50. Avizo Software | Materials Characterization Software - UK [Internet]. [cited 2025 July 15]. Available from: https://www.thermofisher.com/uk/en/home/electron-microscopy/products/software-em-3d-vis/avizo-software.html

51. Consortium TM. Project MONAI [Internet]. Zenodo; 2020 [cited 2025 Oct 28]. Available from: https://zenodo.org/records/4323059

52. Oktay O, Schlemper J, Folgoc LL, Lee M, Heinrich M, Misawa K, et al. Attention U-Net: Learning Where to Look for the Pancreas [Internet]. arXiv; 2018 [cited 2025 Oct 28]. Available from: http://arxiv.org/abs/1804.03999

53. Horton JM, Summers AP. The material properties of acellular bone in a teleost fish. J Exp Biol. 2009 May;212(Pt 9):1413–20.

54. Kolouri S, Park S, Thorpe M, Slepčev D, Rohde GK. Optimal Mass Transport: Signal processing and machine-learning applications. IEEE Signal Process Mag. 2017 July;34(4):43–59.

55. Peyré G, Cuturi M. Computational Optimal Transport: With Applications to Data Science. MAL. 2019 Feb 11;11(5–6):355–607.

56. Jen A. Bright, Bright JA, Emily J. Rayfield, Rayfield EJ. Sensitivity and ex vivo validation of finite element models of the domestic pig cranium. Journal of Anatomy. 2011 Oct 1;219(4):456–71.

57. Zhang L, Cao Z, Zhao Q. Deep learning-aided segmentation combined with finite element analysis reveals a more natural biomechanic of dinosaur fossil. Sci Rep. 2025 Apr 22;15(1):13964.

58. Brunt LH, Roddy KA, Rayfield EJ, Hammond CL. Building Finite Element Models to Investigate Zebrafish Jaw Biomechanics. J Vis Exp. 2016 Dec 3;(118):54811.

59. Taha AA, Hanbury A. Metrics for evaluating 3D medical image segmentation: analysis, selection, and tool. BMC Medical Imaging. 2015 Aug 12;15(1):29.

60. Menze BH, Jakab A, Bauer S, Kalpathy-Cramer J, Farahani K, Kirby J, et al. The Multimodal Brain Tumor Image Segmentation Benchmark (BRATS). IEEE Transactions on Medical Imaging. 2015 Oct;34(10):1993–2024.

61. Salehi SSM, Erdogmus D, Gholipour A. Tversky loss function for image segmentation using 3D fully convolutional deep networks [Internet]. arXiv; 2017 [cited 2025 Oct 28]. Available from: http://arxiv.org/abs/1706.05721

62. Moen E, Bannon D, Kudo T, Graf W, Covert M, Van Valen D. Deep learning for cellular image analysis. Nat Methods. 2019 Dec;16(12):1233–46.

63. Cellpose: a generalist algorithm for cellular segmentation | Springer Nature Experiments [Internet]. [cited 2025 Oct 21]. Available from: https://experiments.springernature.com/articles/10.1038/s41592-020-01018-x?utm_source=chatgpt.com

64. Accelerating segmentation of fossil CT scans through Deep Learning | Scientific Reports [Internet]. [cited 2025 Oct 21]. Available from: https://www.nature.com/articles/s41598-024-71245-1

65. Yu C, Qin F, Li Y, Qin Z, Norell M. CT Segmentation of Dinosaur Fossils by Deep Learning. Front Earth Sci [Internet]. 2022 Jan 27 [cited 2025 Oct 21];9. Available from: https://www.frontiersin.org/journals/earth-science/articles/10.3389/feart.2021.805271/full

66. Yaqoob M, Ishaq M, Ansari MY, Qaiser Y, Hussain R, Rabbani HS, et al. Advancing paleontology: a survey on deep learning methodologies in fossil image analysis. Artif Intell Rev. 2025 Jan 6;58(3):83.

67. Jiang H, Gao M, Li H, Jin R, Miao H, Liu J. Multi-Learner Based Deep Meta-Learning for Few-Shot Medical Image Classification. IEEE J Biomed Health Inform. 2023 Jan;27(1):17–28.

